# Caudate dopamine predicts cognitive effort decisions both in the lab and daily life

**DOI:** 10.64898/2026.09.26.754663

**Authors:** Jennifer L. Crawford, Rachel E. Brough, Sarah A. Eisenstein, Jonathan E. Peelle, Todd S. Braver

**Author notes:** Correspondence should be addressed to: Jennifer Crawford.

## Abstract

Making choices about whether and when to engage cognitive effort is a common feature of human everyday experience with important consequences for academic, career, and health outcomes. Yet, despite its importance, little is understood about the neurochemical mechanisms that underlie individual differences in the willingness to engage in cognitively effortful activities. To examine this question, we employed a multi-modal neuroimaging approach in which participants performed the Cognitive Effort Discounting paradigm (Cog-ED) while simultaneous PET-fMRI was acquired. We assessed both individual differences in dopamine-D2 receptor availability and brain activity during decision-making. In addition, the same participants performed a parametric working memory task (N-Back) and reported the cognitive effort associated with their daily life activities through a seven-day Ecological Momentary Assessment (EMA) protocol. The findings suggest a central role for caudate dopamine-D2 receptors in modulating cognitive effort-related motivation. People with higher caudate dopamine-D2 receptor availability had better working memory function, heightened sensitivity to effort costs, and greater modulation of brain activity during effort valuation. These laboratory findings also translated directly to daily life: higher levels of caudate dopamine-D2 receptors were associated with greater engagement in cognitively effortful activities. Together, the results suggest that how individuals determine the motivational value of cognitively effortful activities strongly depends on the dopamine system.

**Significance Statement:** Weighing the costs and benefits of engaging in cognitive effort is a common feature of daily life. Nevertheless, there is little consensus about how such decisions are implemented in the human brain. This study used a cognitive effort decision-making paradigm during simultaneous PET-fMRI scanning to identify the neurochemical mechanisms engaged while participants decided whether to engage in cognitive effort. Neuroimaging measures were paired with daily life assessments of cognitive effort to directly test if individual differences in dopamine function translate to how people make daily life decisions. The results provide some of the first evidence that dopamine-D2 receptors in the dorsal striatum play an important role in cognitive effort-based decision-making, not only in the laboratory, but also in daily life.

## Introduction

Weighing the costs and benefits of engaging in cognitive effort is central to daily decision-making (1, 2). Much like people are faced with making cost-benefit decisions pertaining to physical effort (e.g., deciding to take the stairs versus the elevator), they must also make decisions between tasks that differ in the degree of cognitive effort required. For example, an employee might need to decide between engaging in a mentally demanding project that could lead to career advancement opportunities versus a less challenging project that would not produce the same opportunities for salary growth. Indeed, such decisions may take on even more importance in our current technological landscape, with the growing temptation to cognitively surrender, and outsource effortful activities to ever more present advances in technology (3). Prior work has shown that individuals vary in their willingness to engage in cognitive effort (4–7). In general, when people are asked to choose between options differing in cognitive effort, they tend to choose the lower effort option (4, 7), especially when the high-effort alternative is very mentally demanding (5). Moreover, there are clear reductions in the motivation to engage in cognitive effort observed across different populations (e.g., aging, Alzheimer’s disease, depression) (1, 8, 9). As such, considerable attention has focused on the significance of identifying core mechanisms that give rise to individual differences in the willingness to engage in cognitive effort and how such differences relate to daily life behavior (10–12). In the current study, we used a multi-modal neuroimaging approach to test the hypothesis that striatal dopamine receptor availability may be a mechanistic source of these individual differences in cognitive effort decision-making.

Across both humans and non-human animals, there is growing evidence that a core set of brain regions is associated with the willingness to engage in cognitive effort. For example, using functional magnetic resonance imaging (fMRI) in humans, researchers found that the subjective value of cognitive effort is encoded by a set of regions, which includes the ventral striatum, ventromedial prefrontal cortex (vmPFC), and dorsal anterior cingulate cortex (dACC) (13). In studies of rodents, both the vmPFC (14) and the striatum (15) were also related to individual differences in cognitive effort-based decision-making. Further support for the centrality of the vmPFC, ventral striatum, and dACC in cognitive effort-based decision-making comes from a recent meta-analysis (16); however, the meta-analysis suggested that canonical valuation regions (e.g., vmPFC) may play a distinct role in decision-making as compared to the ACC (16). Indeed, in a previously published report using the same individuals included in the present study, we found that activity modulation in the dACC reliably scaled with the relative subjective value of cognitive effort and predicted choice on a trial-by-trial basis (17). Studies in rodents also support the role of the ACC in value comparison when animals are asked to weigh the costs and benefits of engaging in cognitive effort (18). Although there are still ongoing debates of the precise neural mechanisms that support the ability of individuals to weigh the costs and benefits of cognitive effort, it is clear that the vmPFC, striatum, and dACC each play a prominent role when making decisions about whether to engage in cognitively effortful activities.

In parallel research, the dopamine system has emerged as a prominent focus of study across both human and non-human studies of effort-based decision-making (19–24). Chemical neuromodulators such as dopamine are viewed as an attractive target of study because they help to adapt the output of fixed brain structures to changes in the environment (21). The dopamine system represents a particularly advantageous focus of investigation because it has dense projections to and from many of the regions that have been identified in studies of effort-based decision-making (e.g., striatum, vmPFC, dACC) (21, 22, 25). One such account of the role of dopamine in effort-based decision-making suggests that mesolimbic dopamine circuits, of which the ventral striatum is a central hub, help to support the willingness to exert either cognitive or physical effort for rewards across both human and non-human animals (13, 26–28). In particular, dopamine signaling in the ventral striatum is thought to bias the cost-benefit tradeoffs of engaging in effort through converging information received from connected cortical regions, such as the ACC (29, 30). Nevertheless, most of the support for this hypothesis comes from studies of physical effort-based decision-making, leaving open the question of whether the same dopaminergic circuits are also involved in the cost-benefit decisions pertaining to cognitive effort.

Indeed, emerging lines of work suggest that it is the dorsal rather than ventral striatum which may be most important in the willingness to engage in cognitive effort. Recent theoretical frameworks have coalesced around the idea that distinct cortico-striatal circuits support different types of cost-benefit decisions (20, 21). Specifically, in this framework, cognitive effort-based decisions are hypothesized to be subserved by nigrostriatal dopamine circuits, which include the substantia nigra, dorsal striatum, and the ACC (20). Recent empirical evidence supports this claim, demonstrating that people with higher dopamine synthesis capacity in the dorsal, but not ventral, striatum tend to choose to engage in cognitive effort more (and have a greater reliance on working memory processes) relative to those with lower dopamine synthesis capacity (31, 32). Similar patterns are observed in the rodent literature, in which the dorsal, but not ventral, striatum was shown to be necessary for trading off the costs and benefits of effortful cognitive actions (33). Together, these results suggest that dopamine function in nigrostriatal circuitry covaries with individual differences in the willingness to engage in cognitive effort.

The current study was aimed to fill an important gap in the current body of work on human cognitive effort-based decision-making, by providing direct evidence regarding the neuromodulatory role of striatal dopamine on both brain activity and behavior during cognitive effort-based decision-making. Namely, although PET imaging can provide evidence of dopamine involvement (i.e., indexing receptor availability and synthesis capacity), this method does not provide evidence regarding whether individual differences in dopamine are related to neural activity metrics of cognitive effort-based decision-making. For that, other neuroimaging modalities are required, such as fMRI. Moreover, there has been a disconnect between work that identifies neural and behavioral markers of cognitive effort decision-making in laboratory tasks and assessments of cognitive effort in daily life contexts. As such, the ecological validity of prior brain and behavioral findings is limited.

We addressed these key issues and limitations by examining the contribution of the striatal dopamine in cognitive effort-based decision-making in both the laboratory and naturalistic contexts. To do so, we combined simultaneous PET-fMRI with Ecological Momentary Assessment (EMA). Specifically, individual differences in dopamine D2-like receptor availability in the striatum [PET] and brain activity modulation [fMRI] were jointly assessed while participants performed the Cognitive Effort Discounting paradigm (Cog-ED) (13, 17), a previously validated neuroeconomic decision-making paradigm, that requires participants to make choices between effortful options systematically varying in subjective value. These measurements were paired with a week-long intensive longitudinal sampling protocol [EMA] that repeatedly assessed daily life cognitive effort (1). To preview, we found clear evidence for the role of dopamine-D2 receptors in the caudate—rather than the ventral striatum—as a reliable predictor of effort-related cognitive capacity, neural encoding of subjective value, cognitive effort-based decision-making, and cognitive effort in daily life.

## Results

### D2-receptor availability is linked to cognitive effort costs through capacity constraints

The Cog-ED paradigm requires participants to make choices about whether to perform tasks that are higher or lower in cognitive effort. In the Cog-ED variant we used, cognitive effort was manipulated in terms of working memory load, and as reflected in different levels of the N-Back paradigm (2-Back, 3-Back, or 4-Back). We tested whether individual differences directly impacted the subjective motivational value of cognitive effort, or instead indirectly through cognitive capacity constraints.

N-Back behavioral performance declined as the level of N increased, in that participants were less accurate, *OR* = 0.56 [0.51, 0.63], *z* =-10.22, *p* < 0.001 (**Figure 1A**), and slower in responding, *B* = 0.04 [0.02, 0.06], *t* =4.10, *p* < 0.001. This pattern replicates many decades of study of the N-Back (34, 35), while also supporting the idea that N-Back working memory load can be used to experimentally manipulate the level of cognitive effort required in a parametric fashion, as we and others have directly established in prior work (1, 4, 36). Consistent with this assumption, in the Cog-ED paradigm, participants subjectively discounted the motivational value of performing the N-Back paradigm as working memory load increased, requiring a higher degree of monetary reward to choose to perform high-load N-Back conditions, relative to a low-load baseline condition. Further, replicating the pattern of results observed in multiple previous studies (1, 4, 5), the more difficult levels of the N-Back were discounted more, or had a lower motivational value, relative to easier task levels, *B* = -0.22 [-0.28, -0.16], *t* =-7.39, *p* < 0.001.

**Figure 1.**
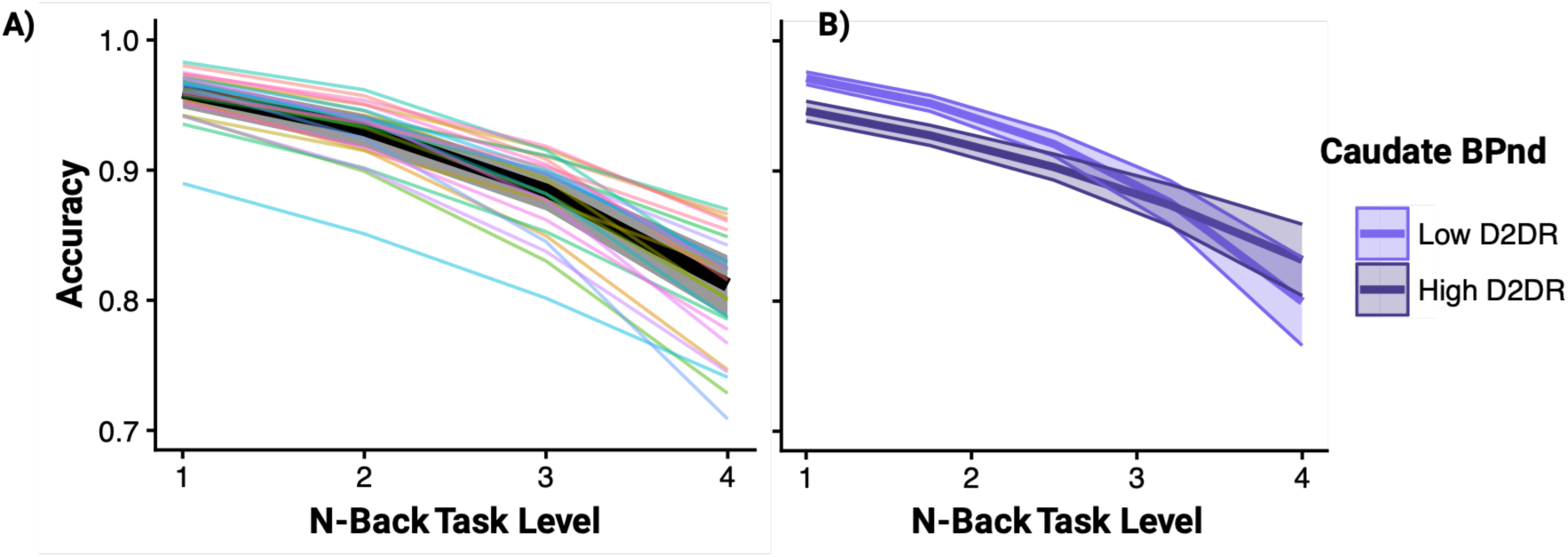
Cognitive performance metrics are associated with striatal dopamine. A) N-Back performance [accuracy] as a function of increasing task load. The black, solid line represents the fixed effect from the generalized linear mixed effect model. The colored lines represent the random effects from the same model (i.e., individual participant’s performance slopes). B) Individual differences in dopamine-D2 receptor availability moderate the effects of task load on N-Back task accuracy. The lines represent ±1SD of caudate dopamine-D2 receptor availability.

Strikingly, we found that the extent to which participants’ accuracy declined across increasing N-Back load levels was meaningfully related to underlying differences in dopamine-D2 receptor availability in the striatum. Namely, dopamine-D2 receptor availability moderated the association between task level and accuracy in the ventral striatum, *OR* = 1.22 [1.05, 1.42], *z* =2.31, *p* = 0.011, and the caudate, *OR* = 1.16 [1.02, 1.32], *z* =2.33, *p* = 0.020 (**Figure 1B**). In other words, people with higher dopamine-D2 receptor availability tended to have less steep of a dropoff in accuracy as task difficulty increased. These effects were selective to accuracy; no such pattern was observed for response times in either striatal region, *ps* > 0.744.

In contrast, however, no effects of dopamine-D2 receptor availability were found on Cog-ED discounting itself. Individual differences in dopamine-D2 receptors were not associated with these baseline estimates of cognitive effort costs in either the ventral striatum, *B* = -0.03 [-0.19, 0.13], *t* =-0.33, *p* = 0.739, or caudate, *B* = 0.04 [-0.08, 0.17], *t* =0.69, *p* = 0.490. Taken together, these results suggest that it is cognitive performance constraints, a frequently observed predictor of cognitive effort costs (4, 5), rather than cognitive effort costs *per se*, which might be the most sensitive individual differences predictor related to dopamine receptor availability in the striatum.

### D2-receptor availability modulates neural correlates of cognitive effort valuation

During fMRI scanning, we assessed neural activity during trials in which participants made repeated decisions in which they chose between performing a high-load N-Back condition, for a higher amount of monetary reward, or a low-load control condition for a lesser reward. Trials were structured so that the first phase presented the high-load / high-reward option in isolation, with both the load-level (2-Back, 3-Back or 4-Back) and monetary reward ($2, $3, or $4) varying randomly across trials (see **Materials and methods**, **Figure S1**), allowing for a pure estimation of the trial-related motivational value of cognitive effort in terms of integrated cost-benefit relationships. We tested whether D2-receptor availability might predict trial-by-trial fluctuations in neural activity associated with encoding the motivational value of high-load cognitive effort.

To test this hypothesis, a priori brain regions of interest were taken from prior neuroimaging studies of the Cog-ED. In this prior work, activity modulation in the dACC, along with the vmPFC and ventral striatum (see **Materials and methods** for more detail) was found to reliably scale with trial-wise estimates of subjective value (13). Across all three regions of interest (ROIs), we found consistent support for the role of caudate dopamine-D2 receptors in subjective value-related activity modulation. Although not significant, subjective value-related activity modulation in the dACC/ACC appeared to have positive associations with individual differences in dopamine-D2 receptor density in the caudate, *B* = 0.13 [-0.02, 0.29], *t* =1.83, *p* = 0.084 (**Figure 2A)**. A more reliable pattern of results was observed in the vmPFC, which is thought to be a key node in the brain’s valuation system ((37); *B* = 0.35 [0.03, 0.68], *t* =2.27, *p* = 0.036; **Figure 2B)**. Even in the ventral striatum, another key node of the valuation system, caudate dopamine-D2 receptor availability was positively associated with subjective value-related activity modulation, *B* = 0.19 [0.02, 0.37], *t* =2.36, *p* = 0.030 (**Figure 2C)**.

**Figure 2.**
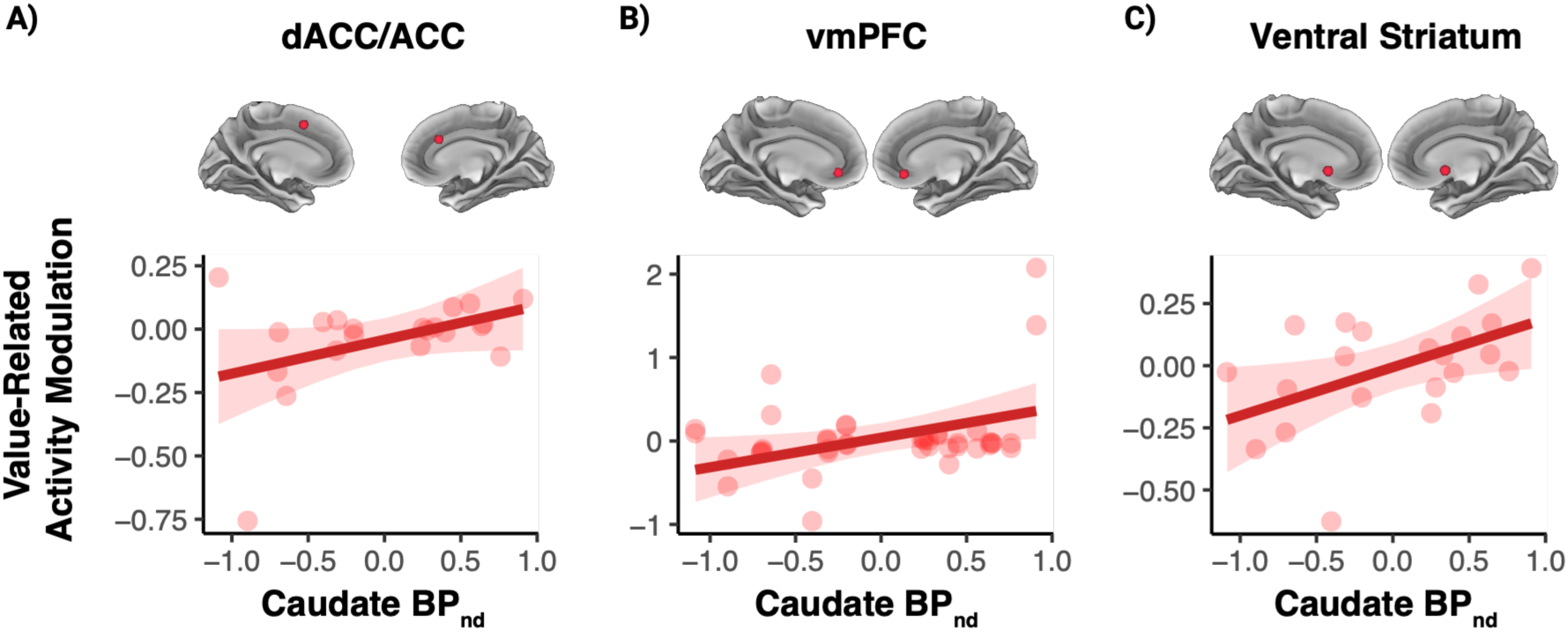
Value-related activity modulation is positively associated with the availability of dopamine-D2 receptors in the caudate. Trial-wise estimates of activity modulation to subjective value in the A) anterior cingulate cortex (ACC) and dorsal ACC (dACC), B) ventromedial prefrontal cortex (vmPFC), and C) ventral striatum covary with underlying individual differences in dopamine-D2 receptor availability in the caudate.

Conversely, in none of these ROIs did dopamine-D2 receptor availability in the ventral striatum predict value-related activity modulation (dACC: *B* = 0.00 [-0.21, 0.22], *t* =0.04, *p* = 0.970; vmPFC: *B* = 0.26 [-0.21, 0.72], *t* =1.15, *p* = 0.266; ventral striatum: *B* = 0.11 [-0.14, 0.36], *t* =0.91, *p* = 0.376). Together, these results suggest that individual differences in dopamine-D2 receptor availability in the caudate, but not in the ventral striatum, are closely associated with neural encoding of motivational value estimates as participants weigh the costs and benefits of engaging in cognitive effort.

### D2-receptor availability modulates cognitive effort-based decision-making

The results above suggest that neural encoding of the subjective motivational value of cognitive effort may be modulated by dopamine-D2 receptor availability in the caudate. We then asked more directly whether caudate D2 receptor availability modulates effort-based decisions themselves, and to what dimensions of the decision are most sensitive to individual variation in this index. Following the valuation phase, in each decision-making trial both the high- and low-effort options were presented together simultaneously in a second phase, and participants chose which they would prefer to perform again for the indicated monetary reward (i.e., participants were told that one of their choices would be selected randomly and they would perform the chosen condition at the end of the session to obtain the offered monetary reward). Choice options were systematically manipulated using a metric that is maximized (+1) when the high effort option has the highest relative subjective value and minimized (-1) when the low effort option has the highest relative value (see **Materials and methods**; **Figure S1**).

Replicating our prior report (17), the relative subjective value of the high-effort option was a reliable predictor of it being chosen, *OR* = 972.04 [188.34, 5016.75], *z* =8.22, *p* < 0.001, independent of the effects of reward, *OR* = 1.56 [1.33, 1.83], *z* =5.43, *p* < 0.001, or task load levels, *OR* = 0.34 [0.29, 0.40], *z* =-13.16, *p* < 0.001. Importantly, the probability of choosing the high-effort choice option was also associated with individual differences in caudate dopamine-D2 receptor availability. Namely, individuals who had higher levels of dopamine-D2 receptors in the caudate had a higher probability of choosing the high effort option (*OR* = 3.79 [1.23, 11.69], *z* =2.32, *p* = 0.020), though this effect was attenuated when controlling for age (*OR* = 2.86 [0.95, 8.62], *z* =1.87, *p* = 0.062). In contrast, we did not observe an association between ventral striatal dopamine-D2 receptors and high-effort choices, *OR* = 2.79 [0.67, 11.67], *z* =1.41, *p* = 0.160.

To more closely investigate relationships between dopamine-D2 receptor availability and choice behavior, we conducted analyses that focused on participants’ sensitivity to the relative costs and benefits of cognitive effort, when deciding between choice options. Specifically, we entered dopamine-D2 receptor availability as a moderating variable between experimentally manipulated parameters reflecting the costs and benefits of engaging in cognitive effort (i.e., relative subjective value, reward level, task load level) and choice behavior. We found that dopamine-D2 receptor availability in the caudate moderated the association between the relative subjective value and choice behavior, *OR* = 5.78 [2.78, 12.05], *z* =4.49, *p* < 0.001 (**Figure 3A)**. In other words, people with higher levels of dopamine-D2 receptors in the caudate were generally more sensitive to shifts in the cost-benefit tradeoff of cognitive effort, being more likely to choose high effort options when the tradeoff was favorable, relative to people with lower receptor availability.

**Figure 3.**
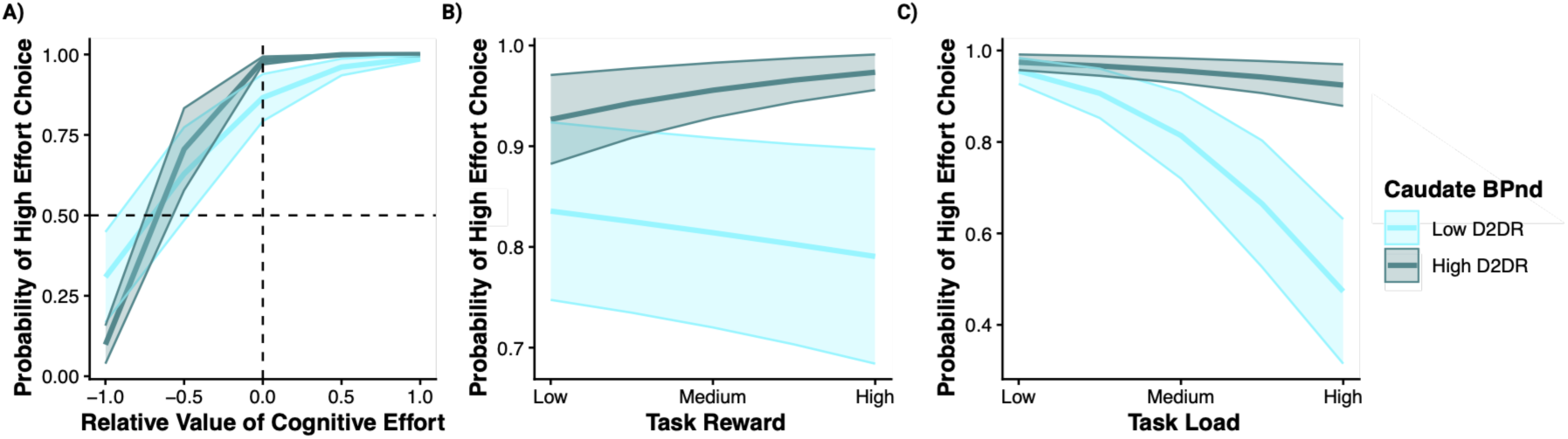
Caudate dopamine moderates the associations between cost and benefits indices and choice behavior during cognitive effort-based decision-making. A) Dopamine-D2 receptor availability moderates the associations between the relative subjective value of cognitive effort and choice behavior. B) The effect of reward level on choice behavior is moderated by caudate dopamine-D2 receptor availability. C) Individual differences in dopamine-D2 receptor availability moderate the effects of task load on choice behavior. Colored lines represent ±1SD of caudate dopamine-D2 receptor availability.

We then examined how these relationships varied according to the constituent elements of relative subjective value, namely effort costs (N-Back task load), and motivational benefits (monetary reward). We observed a moderating effect of caudate dopamine on the relationship between reward level and choice behavior, *OR* = 1.41 [1.06, 1.87], *z* =2.34, *p* = 0.019 (**Figure 3B**). Individuals with low dopamine-D2 receptor availability in the caudate did not strongly modulate choice behavior in relationship to reward level, whereas those with higher levels of dopamine-D2 receptors in the caudate tended to show greater willingness to engage in cognitive effort with increasing levels of reward.

An even stronger moderating effect of caudate dopamine-D2 receptor availability was found on the association between N-Back load level and choice behavior, *OR* = 1.66 [1.23, 2.25], *z* =3.30, *p* = 0.001 (**Figure 3C**). Namely, for individuals with low caudate dopamine-D2 receptor availability, the tendency to choose the high-effort option dropped off steeply across increasing levels of task load, whereas high caudate-D2 individuals tended to choose the high-effort option more frequently and stably across load levels. No such moderating effects were observed for dopamine-D2 receptor availability in the ventral striatum, *ps* > 0.05. Collectively, these findings suggest that caudate dopamine may play an important role in cognitive effort-based decision-making, by modulating sensitivity to the relative costs and benefits associated with cognitive effort engagement.

### D2-receptor availability modulates cognitive effort engagement in daily life

The analyses above explored the relationships between individual differences in striatal dopamine receptors and metrics of cognitive effort measured in controlled laboratory environments, but an important question relates to the ecological validity of such effects. We examined this issue by testing how individual differences in striatal dopamine relate to effort-related indices measured in daily life. In our prior work using the Cog-ED, we showed that people who are more willing to engage in cognitive effort in the laboratory also tend to engage in more cognitively effortful activities in daily life (1). Extending this work, we found that people who engage in more cognitively effortful activities in daily life also have higher levels of dopamine-D2 receptors in the caudate, B = 0.58 [0.18, 0.81], *t* = 2.99, *p* = 0.008 (**Figure 4**). This effect was not observed between estimates of ventral striatal dopamine receptor availability and cognitively effortful daily life activities, B = 0.17 [-0.29, 0.57], *t* = 0.73, *p* = 0.474. Collectively, the pattern of these results suggests that caudate dopamine may serve as central neurochemical mechanism that regulates decisions about whether to engage in cognitively effortful activities, both in the laboratory and in daily life.

**Figure 4.**
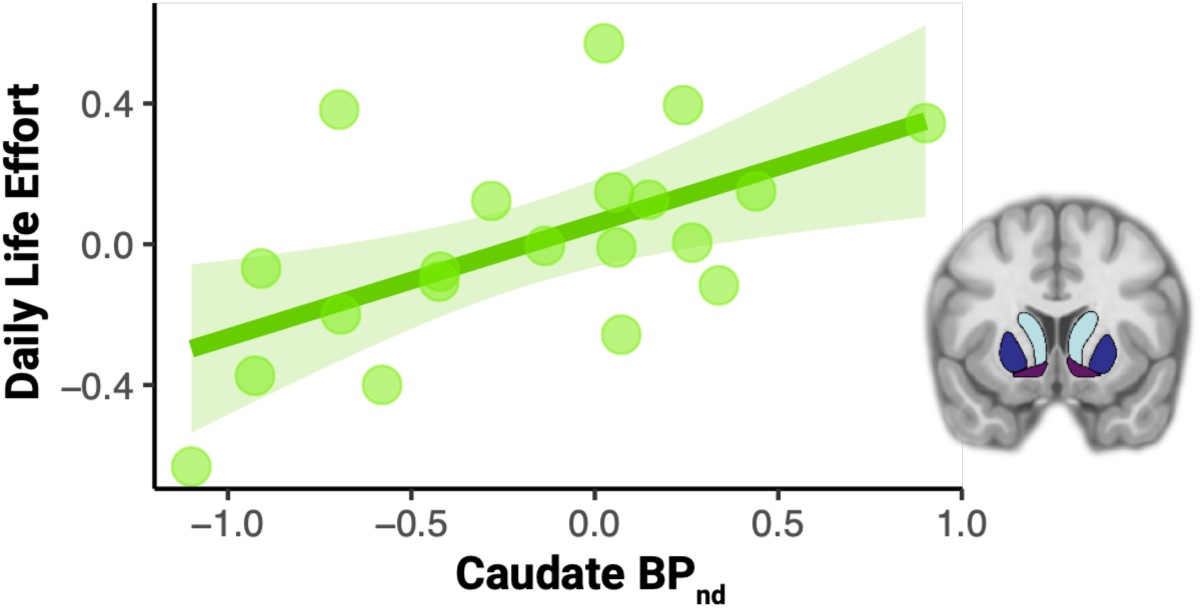
Dopamine in receptor availability in the caudate is positively associated with the mental demand of daily life activities [standardized]. Inset shows striatal regions of interest used in the present study: the caudate (light blue), putamen (dark blue), and ventral striatum (purple).

## Discussion

Decisions involving whether to engage in cognitive effort are central to daily decision-making (1, 2). Dopamine neuromodulation has long been hypothesized as a crucial neurobiological mechanism that helps to explain individual differences in cognitive effort-related motivation (19). Our results provide the first evidence that jointly links together individual differences in the dopamine system with the costs and benefits of engaging in cognitive effort, the neural encoding of cognitive effort motivational value, and engagement in cognitively effortful daily life activities. These findings were made possible by combining simultaneous PET-fMRI with EMA. Put another way, our experimental design enabled measures of dopamine-D2 receptor availability to be linked with both concurrent metrics of behavioral and neural activity assessed during an fMRI task cognitive effort-based decision-making, as well as with naturalistic assessments of cognitive effort engagement during daily-life contexts.

Our results provide clear evidence that dopamine-D2 receptors in the caudate play a central role in modulating cognitive effort-related motivation. These findings represent a significant advance in the growing body of work, across both human and non-human animals, pointing to dopaminergic mechanisms as a central neurobiological substrate of cognitive effort-based decision-making (19–22, 38). In rodent studies, individual differences have also been observed in the willingness to choose to engage in tasks associated with greater cognitive effort. Strikingly, administration of psychostimulants (amphetamine) promoted greater willingness to engage in cognitive effort only for the rodents who originally had low levels of cognitive motivation (39). Studies of non-human primates have also demonstrated that activity in dopamine neurons in the substantia nigra scale with the costs of engaging in cognitive effort (40). In humans, individuals with higher dopamine synthesis capacity in the caudate were tended to engage in cognitive effort with greater frequency than those with lower dopamine synthesis capacity (31). In parallel to the rodent findings, the same study found that administration of a dopamine agonist (methylphenidate) was associated with an increase in the willingness to engage in cognitive effort, but only for people who had low baseline levels of dopamine synthesis capacity in the caudate (31). The results from the current study significantly extend these prior cross-species findings by directly linking individual differences in dopamine receptors to both cognitive effort-based decision-making and effort-related value representations in the brain, while at the same time confirming the ecological validity of such patterns, by demonstrating linkages between dopamine receptors and daily life cognitive effort.

It is notable that our initial pre-registered hypothesis was that dopamine receptor availability in the ventral striatum would be most sensitive to individual differences in cognitive effort-related motivation. This hypothesis was made based on a large body of evidence pointing to dopamine in the ventral striatum as central to value-based decision-making (41, 42). Yet our findings instead revealed a consistent role for dopamine in the dorsal striatum (i.e., caudate) rather than ventral, linking to metrics of cognitive effort engagement both in the laboratory and in daily life. The only other study to examine the linkages between in vivo measures of dopamine and the willingness to engage in cognitive effort in humans also showed that individual differences in dopamine synthesis capacity in the caudate, but not ventral striatum were sensitive to cognitive motivation (31). Work in rodents further implicates this dissociation between the dorsal and ventral striatum in supporting cognitive effort-related motivation; temporary inactivation of the dorsal, but not ventral, striatum decreased the rodents’ preferences for the high-effort, high-reward option and also diminished their performance on the task (15).

Indeed, our findings are highly consistent with the growing recognition that dorsal and ventral subregions of the striatum appear to serve complementary roles in cognitive effort-related motivation (20). Dopamine signaling in distinct subregions is proposed to mediate different types of cost–benefit tradeoffs across hierarchical cortico-striatal circuits, which may help to explain how different aspects of the dopamine system work together when individuals are faced with determining the value of cognitive effort and when to engage in it. In particular, it may be that value-based decisions regarding cognitive effort may be qualitatively distinct from other types of value-based decision-making in that they require estimates of action value rather than purely stimulus or outcome value. It is well-appreciated that dorsal striatum may be more actively involved with decision-making that involves estimating subjective value of different action choices (20, 43, 44). Our work adds to this conceptual framework, and points to the need for further research investigating whether dopamine in the caudate is specifically geared towards the learning and representation of action values needed to make decisions about whether and when it is motivationally advantageous to engage in specific actions, even if these require significant cognitive effort expenditure.

The findings from the current study also offer an important lens through which prior work describing the neural mechanisms underlying cognitive effort-based decision-making can be more precisely understood. For example, in the same set of participants, we previously found that increased dACC activity to the high effort option during the valuation period predicted choice of that option (17). Although speculative, the tone of the dopamine system (e.g., caudate dopamine-D2 receptor availability) could be the source of this effect. In other words, the choice of a high effort option occurs when its relative subjective value is judged to be favorable, relative to a lower-effort alternative. Presumably, people with greater dopamine-D2 receptor availability are more likely to estimate a favorable relative subjective value, all else being equal, and that this is reflected in higher valuation-related activity to that option (e.g., in dACC). This in turn makes the computation of a favorable relative subjective value more likely, leading to a higher likelihood of selecting the higher effort option.

In contrast, our results suggest that people with lower levels of dopamine-D2 receptor availability in the caudate experience high-load N-Back conditions as being more effortful, potentially due to working memory capacity constraints; correspondingly, their performance drops off more precipitously under those conditions. As such, when individuals with low levels of dopamine-D2 receptors in the caudate are estimating the effort costs of those conditions in decision-making situations, they have a higher cost estimate, and lower subjective value of the high-effort option when presented, even when it is associated with high rewards. This could explain why people with lower levels of dopamine-D2 receptors in the caudate are less likely to select the high-effort option when comparing it to a lower-effort one. When viewed collectively, all of these computations are mirrored in cognitive effort choices in daily life, which means that they are less likely to engage in mentally effortful activities as they go about their daily lives. Extending beyond the healthy, young adult population characterized in the present work, our findings point to caudate dopamine as a promising target of investigation for groups of people with known deficits in cognitive effort-related motivation. For example, our prior work suggests that the Cog-ED may be a particularly useful tool for investigating the neurobiological substrates underlying cognitive effort-based decision-making in older adults, as we have found consistent evidence for reduced motivation to engage in cognitive effort within this population (1, 8). The present study suggests the possibility that such effects might result from age-related declines in dopamine-D2 receptors in the caudate. Indeed, prior work has already shown that dopamine-D2 receptors in the caudate tend to decline with age (45, 46). Measures of the dopamine system have also exhibited linkages with N-Back performance and decision making in older adults (47, 48). As such, future work that combines behavioral and fMRI measures of cognitive effort discounting with PET measurements of the dopamine system will be best suited to disentangle the relationships across these variables and quantify the impact of age-related changes on dopamine function and cognitive effort-based decision-making.

The current study had several strengths, including the use of a multimethod approach involving both simultaneous PET-fMRI and daily life sampling; nonetheless, there were also some notable limitations that point to important future research directions. First and foremost, although this is one the of largest studies to use simultaneous PET-fMRI involving dopamine receptor imaging and measures of cognitive task BOLD signals, our sample size was still limited for an individual differences-focused study. As a consequence, we did not have sufficient statistical power to conduct formal tests of caudate dopamine as a mediator of key individual differences effects. Although simultaneous PET-fMRI imaging studies are both quite technically challenging and expensive to conduct, our work points to the need for the investment in a follow-up study with a large enough sample size to more rigorously test causal hypotheses regarding the role of caudate dopamine in mediating the relationship between cognitive effort exertion (e.g., during tasks such as the N-Back), effort discounting during value-based decision-making, and daily-life effort engagement. Similarly, because of limited statistical power, our study used ROI-focused analyses, utilizing regions defined in prior work with the Cog-ED, and consequently may have missed identifying other regions that could be revealed from whole-brain analyses. Finally, we focused here on the use of NMB as a selective and non-displaceable radiotracer for dopamine D2-like receptors. Although NMB is highly sensitive to dopamine-D2 receptor availability in the striatum, it is not as well-suited to quantify extrastriatal dopamine receptors, particularly cortical dopamine receptors, relative to other radiotracers (e.g., FLB457-D2/D3 (49); NNC112-D1/D5 (50)) and is also insensitive to endogenous displacement of dopamine, such as under task conditions (cf. raclopride) (51). Thus, a priority for future work would be to replicate this paradigm with other radiotracers that can reliably assess the dopamine system.

Nevertheless, the present study provides some of the first data demonstrating a role for dopamine-D2 receptor availability in the caudate for human cognitive effort-based decision-making across both the laboratory and daily life. Crucially, caudate dopamine-D2 receptors predicted the likelihood of an individual choosing to engage in a task associated with higher cognitive effort, brain activity modulation during effort valuation, effort in daily life, and cognitive constraints associated with the willingness to engage in cognitive effort. As such, these findings provide a clear foundation for further work aimed at investigating the potential role of the dopamine system in promoting cognitively effortful behaviors in both health and disease, and across the lifespan, particularly in populations for which apathy is a clinical marker (e.g., pathological aging).

## Materials and Methods

### Participants

Participants were adults, ages 18-40, recruited through Washington University School of Medicine subject recruitment registries. Study participation was subject to the following exclusion criteria: any history of head trauma, any significant medical condition, or any condition that would interfere with MRI or PET imaging (e.g., inability to fit in the scanner, claustrophobia, cochlear implant, metal fragments in eyes, cardiac pacemaker, neural stimulator, pregnancy, metallic body inclusions or other contraindicated metal implanted in the body, or routine occupational exposure to radiation). Participants were also excluded if they reported a history of substance abuse, current tobacco use, alcohol consumption greater than eight ounces of whiskey (or equivalent) per week, use of psychostimulants (excluding caffeine) more than twice at any time in their life (or at all in the past six months), or the use of any psychotropic medication in the last six months. All pre-menopausal women had negative pregnancy tests on the day of the scan. A total of twenty-six participants completed the study, however five of these participants did not have simultaneously acquired PET data because of quality control failures or inability to synthesize the radiotracer on the day of the scan. As such, the final sample consisted of 21 participants (*M_age_*=27.9, *SD_age_*=5.7; 15 females; 3 Asian, 4 Black or African American, 12 White, 2 more than one race; 2 Hispanic or Latinx). All experimental procedures were approved by the Washington University Human Research Protections Office prior to data collection. All participants provided informed consent and were compensated $50/hour for study procedures.

### Design

#### Behavioral Visit

Participants first came into the lab to complete the Cognitive Effort Discounting paradigm (Cog-ED) (4). This enabled participants to become familiar with the tasks that they would later perform in the scanner as well as providing a baseline measure of each participant’s working memory ability and cognitive effort costs (to be used in the neuroimaging visit). As a brief overview, in the Cog-ED, participants were first familiarized with the cognitive effort required to perform different conditions of a demanding working memory task (i.e., N-Back). In the N-Back, working memory load was varied across task levels (N = 1-4). Participants were asked to indicate when the current stimulus (i.e., letter) matched the letter from N steps earlier in the sequence (target) or when the stimulus differed from the letter presented N steps earlier (non-target), with higher levels of N indicating increased cognitive demands. Each task run contained sixty-four trials; 25% of the trials (i.e., 16 trials) were targets. Participants performed two runs of each level of N in ascending order, for a total of 528 trials. On each trial, accuracy and response time were measured. Participants were also asked to rate their perceived levels of mental demand, effort, frustration, and task performance after each level of the task using the NASA Task Load Index (52).

Next, participants completed the subjective value estimation phase of the task. Namely, in the Cog-ED, participants make a series of decisions between performing high-effort task levels (i.e., 2-4 back) for a fixed, high monetary reward ($2, $3, or $4) or low-effort task levels (i.e., 1-back) for a lower monetary reward value. As participants make decisions, the offer for the smaller reward is stepwise titrated until participants are indifferent between the two offers (i.e., they would choose either offer equally). These indifference points estimate the “cost” of cognitive effort. From the indifference point, we compute the subjective value; this value is calculated by dividing the indifference point by the value of the high-effort choice option. In other words, the subjective value estimates reflect the proportion of value an individual is willing to forgo to avoid performing the more effortful task.

#### Neuroimaging Visit

During scanning, participants performed make repeated decisions in a trial-based decision-making variant of the Cog-ED, which was adapted for neuroimaging contexts with a previously published design (13, 17). Each trial began with a valuation phase, during which only the high-effort/high-reward option was presented for six seconds, allowing estimation of its neural encoding (see **Figure S1**). As such, participants were instructed to take this time and consider what the offer was worth to them.

Following the valuation phase of the trial, participants chose between either a low- or high-effort offering with an associated monetary reward. In this decision-making phase, the low- and high-effort options were systematically manipulated according to individually estimated indifference points (obtained from the behavioral visit). Specifically, we manipulated the dollar amount of the low-effort option in relation to the subjective value of the high-effort/high-reward option presented on that trial. This scaling parameter ranged from -0.4 to -0.6 and +0.1 to +0.2, in addition to a subset of trials (N= 18), which had scaling parameters equal to either -1 or +1 (i.e., “catch” trials). Put another way, on a trial in which the scaling parameter was equal to +0.2, the subjective value of the low-effort option is 20% below the subjective value of the high-effort option, which should bias the participant to choose the high-effort/high-reward option (**Figure S1**). Participants completed three runs of the Cog-ED for a total of 108 trials (36 trials/run). All responses were made with a button box provided to participants in the scanner.

#### Daily Life Sampling

To examine the relationships between individual differences in dopamine receptor availability and the cognitive effort of activities in daily life, we asked participants to complete a previously established Ecological Momentary Assessment (EMA) protocol (1). After completing the initial behavioral session in the lab, participants completed a tutorial that walked them through the necessary steps to download the EMA app (Expiwell; https://www.expiwell.com) and complete the EMA surveys. First, participants were instructed to download Expiwell on their phone. Next, participants were taken through the structure and content of the EMA survey (including all survey items) and were asked to answer questions about the contents of the survey to ensure proper understanding before beginning the protocol. As a part of the 7-day EMA protocol, participants received five randomly prompted surveys per day over a fixed twelve-hour window, with approximately two hours in between each survey, for a total of 35 assessment points.

For each survey, participants received a notification containing a link to complete the survey through Qualtrics (Qualtrics, Provo, UT). Upon clicking on the survey link, they were asked to record the activities they engaged in during the past two hours from a list of activities. Following this prompt, a subset of up to three of these previously endorsed activities were randomly sampled for participants to answer an additional set of questions. Specifically, these questions asked participants to rate how mentally demanding each of these activities was (using a rating scale ranging from “Not at all” (1) to “Extremely” (5)) and their level of motivation and enjoyment toward each activity. Directly relevant to this study, the survey items indexing daily life activities and mental demand were taken from previous work, which demonstrated that people who are more willing to engage in cognitive effort in the lab (i.e., those who have lower cognitive effort costs) tend to have higher mental demand of daily life activities (1). Participants were allotted 30 minutes to begin the survey after each notification from the app, after which time their responses were not recorded. If participants did not complete the survey within fifteen minutes after the initial notification, they automatically received a reminder to complete the survey. Overall, participants completed 75.7% of all surveys.

### Data Acquisition and Preprocessing

Scanning was performed on a Siemens 3T Biograph mMR, capable of simultaneous PET-MRI acquisition. With PET, we measured dopamine D2-receptor binding using the highly selective and non-displaceable radioligand N-[C-11]- (methyl)benperidol(C11-NMB)(53), which unlike many other PET tracers shows specific affinity in dopamine-D2, but not D3 receptor availability. As such the C11-NMB radioligand data can provide an index of stable individual differences in dopamine D2-receptor availability in striatal brain regions. Acquisition of the PET data was synchronized to the onset of fMRI scanning. After a bolus intravenous injection of C11-NMB (10-30 mCi; sample mean = 18.77 mCi), dynamic PET emission data were collected in list mode over a 120-minute period (3 x 1-min, 4 x 2-min, 3 x 3-min, and 20 x 5-minute frames).

Preprocessing and analysis of PET data was implemented with PMOD software. After MR-based attenuation correction, PET scan frames were corrected for motion with the 15th dynamic image frame of the series serving as the reference image; a 6 mm FWHM kernel was used to smooth the data. The realigned PET frames were then merged and reassociated with their acquisition timing info in PMOD’s PVIEW module to create a single 4D file for use in PMOD’s PNEURO tool for further analysis. Time activity curves (TACs) from each region were extracted from the PET data and fit with a Logan graphical model (54, 55), which uses the last sixty minutes of the data for the kinetic modeling. A bilateral gray matter cerebellum ROI was used as the reference region using PMOD’s PKIN module for these analyses. Striatal ROIs (caudate, putamen, ventral striatum) were defined using the deep nuclei atlas in PNEURO. Within each participant, mean non-displaceable binding potential (BP_nd_) was calculated for each ROI in each hemisphere. A descriptive summary of all BP_ND_ estimates can be found in the Supplement (**Table S1**, **Figure S3**).

MRI data were acquired and analyzed following standardized procedures, which included high-resolution anatomical (MPRAGE, 0.8mm^3^ voxels) and whole-brain BOLD EPI scans (32-channel head-coil, 4mm^3^ voxels, 32 slices, TR=2000 msec). Preprocessing for both anatomical and functional MRI data was performed using fMRIPrep (56). fMRIPrep performs basic processing steps, such as coregistration, normalization, unwarping, noise component extraction, segmentation, skullstripping etc (see **Supplement** for the detailed fMRIprep processing information). The preprocessed BOLD runs were smoothed with an 8 mm FWHM kernel using 3dBlurtoFWHM from AFNI (57), as well as scaling using 3dTstat and 3dcalc from AFNI. Finally, the images were reoriented to LPI orientation using AFNI’s 3dresample function.

General linear models (GLMs) were applied to extract beta estimates for the event-related activity for each participant. Specifically, AFNI’s 3dDeconvolve function was used to set up the GLMs. In the analyses described below, we used amplitude-modulated tent functions time locked to the onset of each trial. In other words, we quantified the activity modulation associated with trial wise changes in the value of the high-effort, high-reward choice option (using participant’s subjective value estimates measured in the behavioral session) during the valuation phase of the trial (5-9 s after trial onset). The GLMs were run with 3dREMLfit from AFNI. A regressor for the six motion parameters generated from fMRIPrep was included in the GLMs as nuisance regressors. Additionally, TRs were censored if the derivative values were estimated to have a Euclidian norm > 0.3mm. Next, these beta estimates were extracted for each region of interest (ROI). ROIs were defined using 6mm spheres centered around regions shown to encode subjective value in prior work(13). Specifically, these ROIs included the ventral striatum (MNI_xyz_: Left = -12,12,-6; Right = 12,10,-6), ventromedial prefrontal cortex (vmPFC) (MNI_xyz_: Left = -7,38,-11; Right = 4,35,-12), anterior cingulate cortex (ACC) (MNI_xyz_ = -2,28,28), and dorsal ACC (MNI_xyz_ = -2,16,46). All beta estimates were averaged for all voxels within each ROI using 3dROIstats from AFNI. To limit the number of comparisons, across all analyses the values from the left and right hemisphere values were averaged together; values from ACC and dACC were also averaged together.

### Quantification and Statistical Analysis

Data analysis was conducted in R version 4.4.2 (58) using the ‘easystats’ ecosystem (59–63). All multilevel models were conducted with the package *lme4* (version 1.1-36)(64). Across all models, effects are reported as the estimate from the model with 95% confidence intervals and the corresponding test statistic and p-value. All data and analysis code used for the present study can be found on the Open Science Framework: https://osf.io/4v68b.

#### Behavioral Data Analysis

Repeated measures of N-Back task performance and Cog-ED decision-making were characterized using linear mixed effects models. N-Back accuracy was modeled with a generalized linear mixed effect model; task level was modeled as both a fixed and random effect and random effects estimates were extracted to test for their relationship with dopamine-D2 receptor availability. Cog-ED choice behavior was also modeled using a generalized linear mixed effect model, with fixed effects of task load, reward amount, and relative subjective value; relative subjective value was also modeled as a random effect. Random effect estimates from this model were used in subsequent analyses that examined its associations with measures of striatal dopamine.

#### Neuroimaging Data Analysis

We used linear models to characterize the relationships between dopamine-D2 receptor availability in the striatum with task behavior, brain activity modulation, and daily life effort. Because we observed significant age effects in our striatal ROIs (see **Supplement**; **Figure S3**), all analyses using striatal binding potential estimates included age as a covariate, unless otherwise noted. Separate models were used to test the influence of dopamine-D2 receptors in the caudate vs. ventral striatum. In each model binding potential estimates and age were defined as independent variables, effort-related variables were defined as the dependent variable across all hypothesis tests.

## Acknowledgments

The research was supported by the National Institutes Health (NIH; R21 AG067295 to T.S.B. and subaward R24 AG054355 to J.L.C.). J.L.C. was additionally supported by T32 AG000030 and F32 AG085890. As such, this work is subject to the NIH Public Access Policy. Through acceptance of this federal funding, NIH has been given a right to make this manuscript publicly available in PubMed Central upon the Official Date of Publication, as defined by NIH.

Rachel Brough is now affiliated with the Department of Psychology at the University of Denver. Sarah Eisenstein is now affiliated with the Brown School of Social Work at Washington University. Jonathan Peelle is now affiliated with the Departments of Communication Sciences and Disorders and Psychology at Northeastern University.

## Author Contributions

J.L.C., S.A.E, J.E.P., and T.S.B. designed the research; J.L.C and R.E.B performed the research; J.L.C. analyzed the data, prepared data visualizations, and wrote first draft of the manuscript; R.E.B., S.A.E., J.E.P., and T.S.B. edited the manuscript.

