## Supplemental material for "Caudate dopamine predicts cognitive effort decisions both in the lab and daily life"

**Supplemental Materials**

The following document contains supplemental information and figures that accompany the manuscript, *Caudate dopamine predicts cognitive effort decisions both in the lab and daily life*.

**Cognitive Effort Discounting Task**
During simultaneous PET-fMRI neuroimaging participants are first presented with a valuation phase (six seconds) in which only the high-effort, high-reward option is shown.

Following this, participants begin the decision phase of the trial (six seconds) in which two options are presented to them, and they are instructed to choose the option they prefer. Importantly, the subjective value (SV) of the low-effort, low-reward option is systematically manipulated on each trial to bias participants to choose the low- ("easy" = -1; "hard" = - 0.2 to -0.1) and high-effort options ("hard" = 0.4 to 0.6; "easy" = 1) equally.

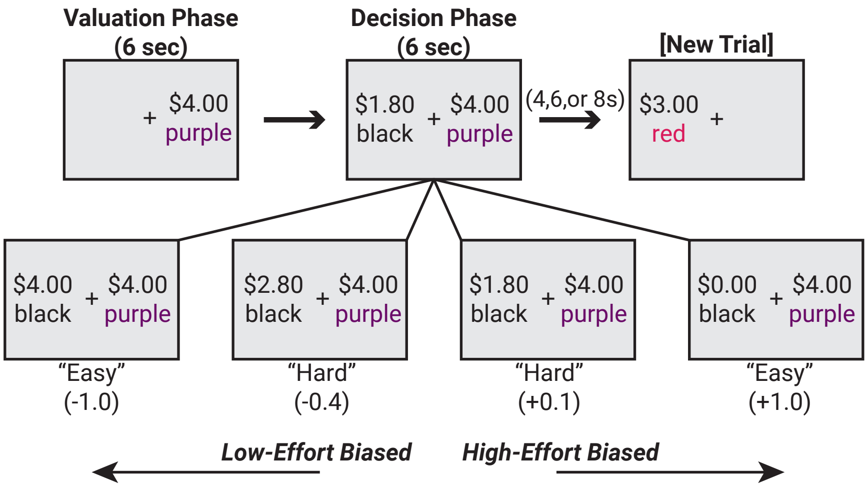


**Figure S1.** Cog-ED task schematic.

**fMRIPrep Details**

The T1-weighted (T1w) image was corrected for intensity non-uniformity (INU) with N4BiasFieldCorrection [1], distributed with ANTs 2.3.3 [2]; RRID:SCR_004757), and was used as T1w-reference throughout the workflow. The T1w-reference was then skull-stripped with a *Nipype* implementation of the antsBrainExtraction.sh workflow (from ANTs), using OASIS30ANTs as target template. Brain tissue segmentation of cerebrospinal fluid (CSF), white-matter (WM) and gray-matter (GM) was performed on the brain-extracted T1w using fast (FSL 5.0.9, RRID:SCR_002823; [3]. Volume-based spatial normalization to one standard space (MNI152NLin2009cAsym) was performed through nonlinear registration with antsRegistration (ANTs 2.3.3), using brain-extracted versions of both T1w reference and the

T1w template. The following template was selected for spatial normalization: *ICBM 152 Nonlinear Asymmetrical template version 2009c* [[4]; RRID:SCR_008796; TemplateFlow ID: MNI152NLin2009cAsym].

For each of the BOLD runs per participant (across all tasks), the following preprocessing was performed. First, a reference volume and its skull-stripped version were generated using a custom methodology of *fMRIPrep*. A B0-nonuniformity map (or *fieldmap*) was estimated based on a phase-difference map calculated with a dual-echo GRE (gradient-recall echo) sequence, processed with a custom workflow of *SDCFlows* inspired by the epidewarp.fsl script and further improvements in HCP Pipelines [5]. The *fieldmap* was then co-registered to the target EPI (echo-planar imaging) reference run and converted to a displacements field map (amenable to registration tools such as ANTs) with FSL's fugue and other *SDCflows* tools. Based on the estimated susceptibility distortion, a corrected EPI (echo-planar imaging) reference was calculated for a more accurate co-registration with the anatomical reference. The BOLD reference was then co-registered to the T1w reference using flirt (FSL 5.0.9;[6] with the boundary-based registration[7] cost-function. Co-registration was configured with nine degrees of freedom to account for distortions remaining in the BOLD reference. Head-motion parameters with respect to the BOLD reference (transformation matrices, and six corresponding rotation and translation parameters) are estimated before any spatiotemporal filtering using mcflirt (FSL 5.0.9; [8]. BOLD runs were slice-time corrected using 3dTshift from AFNI 20160207 [9]; RRID:SCR_005927).

The BOLD time-series (including slice-timing correction when applied) were resampled onto each subject's native space by applying a single, composite transform to correct for head-motion and susceptibility distortions. These resampled BOLD time-series will be referred to as *preprocessed BOLD in original space*, or just *preprocessed BOLD*. The BOLD time-series were resampled into standard space, generating a *preprocessed BOLD run in MNI152NLin2009cAsym space*. First, a reference volume and its skull-stripped version were generated using a custom methodology of *fMRIPrep*. Several confounding time-series were calculated based on the *preprocessed BOLD*: framewise displacement (FD), DVARS and three region-wise global signals. FD was computed using two formulations following Power (absolute sum of relative motions)[10] and Jenkinson (relative root mean square displacement between affines)[8]. FD and DVARS are calculated for each functional run, both using their implementations in *Nipype* (following the definitions by Power et al., 2014). The three global signals are extracted within the CSF, the WM, and the whole-brain masks. Additionally, a set of physiological regressors were extracted to allow for component-based noise correction (*CompCor*)[11].

Principal components are estimated after high-pass filtering the *preprocessed BOLD* time-series (using a discrete cosine filter with 128 s cut-off) for the two *CompCor* variants: temporal (tCompCor) and anatomical (aCompCor). tCompCor components are then calculated from the top 2% variable voxels within the brain mask. For aCompCor, three probabilistic masks (CSF, WM and combined CSF+WM) are generated in anatomical space. The implementation differs from that of Behzadi et al. in that instead of eroding the masks by 2 pixels on BOLD space, the aCompCor masks are subtracted a mask of pixels that likely contain a volume fraction of GM. This mask is obtained by thresholding the corresponding partial volume map at 0.05, and it ensures components are not extracted from voxels containing a minimal fraction of GM. Finally, these masks are resampled into BOLD space and binarized by thresholding at 0.99 (as in the original implementation). Components are also calculated separately within the WM and CSF masks. For each CompCor decomposition, the *k* components with the largest singular values are retained, such that the retained components' time series are sufficient to explain 50 percent of variance across the nuisance mask (CSF, WM, combined, or temporal). The remaining components are dropped from consideration. The head-motion estimates calculated in the correction step were also placed within the corresponding confounds file.

The confound time series derived from head motion estimates and global signals were expanded with the inclusion of temporal derivatives and quadratic terms for each[12]. Frames that exceeded a threshold of 0.5 mm FD or 1.5 standardized DVARS were considered motion outliers. All resampling can be performed with *a single interpolation step* by composing all the pertinent transformations (i.e., head-motion transform matrices, susceptibility distortion correction when available, and co-registrations to anatomical and output spaces). Gridded (volumetric) resampling was performed using antsApplyTransforms (ANTs), configured with Lanczos interpolation to minimize the smoothing effects of other kernels[13]. Non-gridded (surface) resampling was performed using mri_vol2surf (FreeSurfer). The preprocessed BOLD runs were smoothed with an 8 mm FWHM kernel using 3dBlurtoFWHM from AFNI [14], as well as scaling using 3dTstat and 3dcalc from AFNI. Finally, the images were reoriented to LPI orientation using AFNI's 3dresample function.

Many internal operations of *fMRIPrep* use *Nilearn* 0.6.2 [15](RRID:SCR_001362), mostly within the functional processing workflow. For more details of the pipeline, see the section corresponding to workflows in *fMRIPrep*'s documentation.

**Descriptive information: striatal binding potential estimates**

Binding potential estimates ($BP_{ND}$) were quantified using PMOD. Below, we detail the descriptive information pertaining to these estimates.

| Region | N | $BP_{nd}$ | SD | SE | CI |
|---|---|---|---|---|---|
| Caudate (left) | 21 | 2.821468 | 1.316653 | 0.2873172 | 0.5993331 |
| Caudate (right) | 21 | 2.658967 | 1.423435 | 0.3106189 | 0.6479396 |
| Putamen (left) | 21 | 3.981932 | 1.402932 | 0.3061449 | 0.6386071 |
| Putamen (right) | 21 | 3.859982 | 1.380854 | 0.301327 | 0.628557 |
| Ventral striatum (left) | 21 | 2.8994 | 1.226584 | 0.2676627 | 0.5583345 |
| Ventral striatum (right) | 21 | 3.060872 | 1.256098 | 0.2741031 | 0.5717691 |

**Table S1.** Binding potential estimates across striatal ROIs.

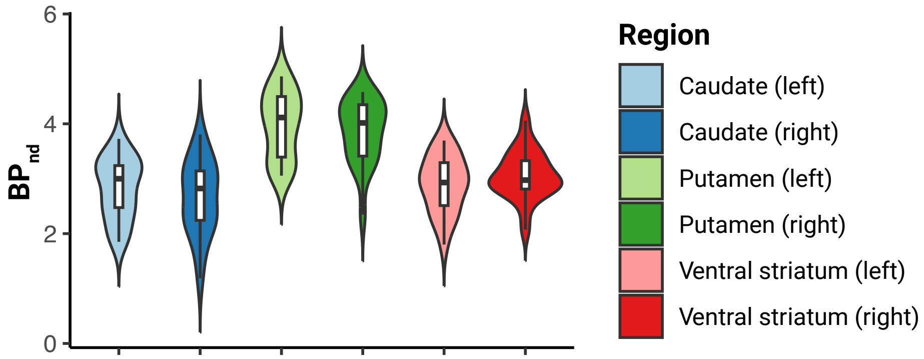


**Figure S2.** Binding potential estimates across striatal ROIs.

Binding potential estimates also covaried with age, $B$ = -0.03 [-0.05, -0.01], $t$ =-2.26, $p$ = 0.025 (**Figure S3)**. As such, age was included as a covariate in all analyses, unless otherwise noted.

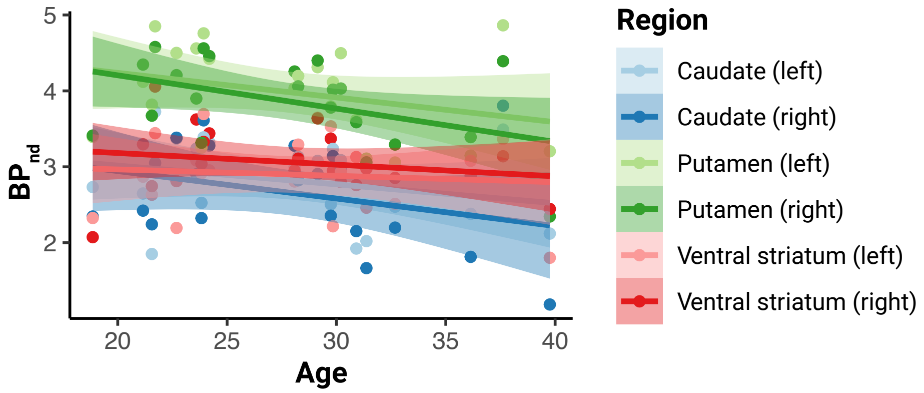


**Figure S3.** Binding potential estimates covary with age.
